# ARID1A regulates R-loop associated DNA replication stress

**DOI:** 10.1101/2020.11.16.384446

**Authors:** Shuhe Tsai, Emily Yun-chia Chang, Louis-Alexandre Fournier, James P. Wells, Sean W. Minaker, Yi Dan Zhu, Alan Ying-Hsu Wang, Yemin Wang, David G. Huntsman, Peter C. Stirling

**Affiliations:** Terry Fox Laboratory, BC Cancer, Vancouver Canada; Department of Molecular Oncology, BC Cancer, Vancouver Canada; Department of Medical Genetics, University of British Columbia, Vancouver Canada; Department of Interdisciplinary Oncology, University of British Columbia, Vancouver Canada

**Keywords:** ARID1A, R-loops, replication stress, BAF complex, Topoisomerase II

## Abstract

ARID1A is lost in up to 7% of all cancers, and this frequency increases in certain cancer types, such as clear cell ovarian carcinoma where ARID1A protein is lost in about 50% of cases. While the impact of ARID1A loss on the function of the BAF chromatin remodeller complexes is likely to drive oncogenic gene expression programs in specific contexts, ARID1A also binds genome stability regulators such as ATR and TOP2. Here we show that ARID1A loss leads to DNA replication stress associated with R-loops and transcription-replication conflicts in human cells. These effects correlate with altered transcription and replication dynamics in ARID1A knockout cells and to reduced TOP2A binding at R-loop sites. Together this work extends mechanisms of replication stress in ARID1A deficient cells with implications for targeting ARID1A deficient cancers.

## INTRODUCTION

Dysregulation of DNA replication and transcriptional programs in cancer cells is likely to give rise to increased transcription-replication conflicts (TRCs) [1]. Such conflicts can create DNA damage and promote mutagenesis. Mutations in key tumour suppressor genes and activation of oncogenes have both been linked to increases in TRCs. These include loss of P53, loss of fork protection and homologous recombination proteins (e.g. BRCA1, BRCA2, Fanconi Anemia factors), activation of H-RAS, or various oncogenic fusion proteins such as EWS-FLI1 or SS18-SSX [2–7]. In most cases these increases in TRCs are associated with persistence or stabilization of an R-loop structure in the genome. R-loops refer to RNA reannealed to a genomic template to create an RNA:DNA hybrid surrounded by a single-stranded loop of the non-template DNA strand [8]. R-loops serve normal regulatory and topological roles in the genome, but their persistence and association with TRCs make them important contributors to genome instability [9]. Given the ever-expanding network of genetic factors that can lead to R-loops and TRCs, additional insights are required to understand the circumstances in which TRCs are damaging, and potentially exploitable for therapeutic benefit.

The BAF (BRG1/BRM-associated factor) chromatin remodeling complex is a frequent target of mutagenesis in cancer. BAF is the human orthologue of the yeast SWI/SNF complex and assembles from between 14-16 subunits in human cells [10]. In general, the BAF complex maintains open chromatin at enhancer elements to drive gene expression programs in a developmentally regulated and cell context specific manner. In addition to regulating gene expression, the BAF complex controls various other aspects of chromatin topology, including notable functions in DNA repair [10–12]. Through interactions with BRIT/MCPH1 BAF is recruited to DNA double strand breaks and plays roles in regulating DNA end resection[12], although its relative contributions to homologous recombination versus non-homologous end joining is yet to be resolved [13–15].

The DNA binding subunit ARID1A, and the catalytic subunit SMARCA4 (BRG1) are the two most mutated BAF subunits across cancer types. ARID1A generally acts as a tumour suppressor gene which is lost in as many as 7% of all cancers, including at very high frequency in certain ovarian cancers (e.g. 50% in clear cell ovarian cancers)[16]. Interestingly, ARID1A may also behave as an oncogene in some settings such as in early lesions of hepatocellular carcinoma[17]. Cells lacking ARID1A can still form an alternative BAF complex incorporating a paralogous protein called ARID1B. Indeed, conditional ARID1A knockout shows that BAF genome occupancy rapidly recovers through the ARID1B-containing complex[18]. Some, but not all, ARID1A functions can be compensated by ARID1B.

Here using isogenic cell lines we show that loss of ARID1A is associated with increases in R-loop abundance, despite overall decreases in global transcription. These excess R-loops are associated with replication stress, DNA breaks, and TRCs, while R-loop suppression restores genome integrity in ARID1A deficient cells. We show that failure to target TOP2A to R-loop prone genomic loci is a mechanism underlying at least part of the increase in R-loop associated TRCs and DNA damage. Together these data define a function for the BAF complex in suppressing TRCs.

## RESULTS AND DISCUSSION

### Transcription-replication conflicts and R-loops accumulate in ARID1A knockout cells

Given recent reports that ATR inhibition is selectively toxic to ARID1A-deficient cells in some settings [19], we tested whether ARID1A knockout elicits replication stress in an isogenic cell line model of clear cell ovarian cancer. We used CRISPR to remove ARID1A (**Supplemental Figure S1A**), and then examined the phosphorylation of RPA2 on serine 33, a known mark of replication stress, and found higher levels of RPA2-S33P by immunofluorescence in ARID1A-deficient cells (**Figure 1A**). To further support the presence of ongoing replication stress we used SIRF (in <u>s</u>itu analysis of protein <u>i</u>nteractions at DNA <u>r</u>eplication <u>f</u>orks) to detect recruitment of Mre11 to nascent EdU labeled DNA [20]. This analysis showed a significant increase in Mre11-SIRF signals in ARID1A-deficient compared to proficient cells (**Figure 1B**), which we and others have previously shown is a symptom of replication stress [21,22]. Together these data suggest that ARID1A knockout cells experience increased replication stress, which recruits MRE11 to nascent DNA.

**Figure 1.**
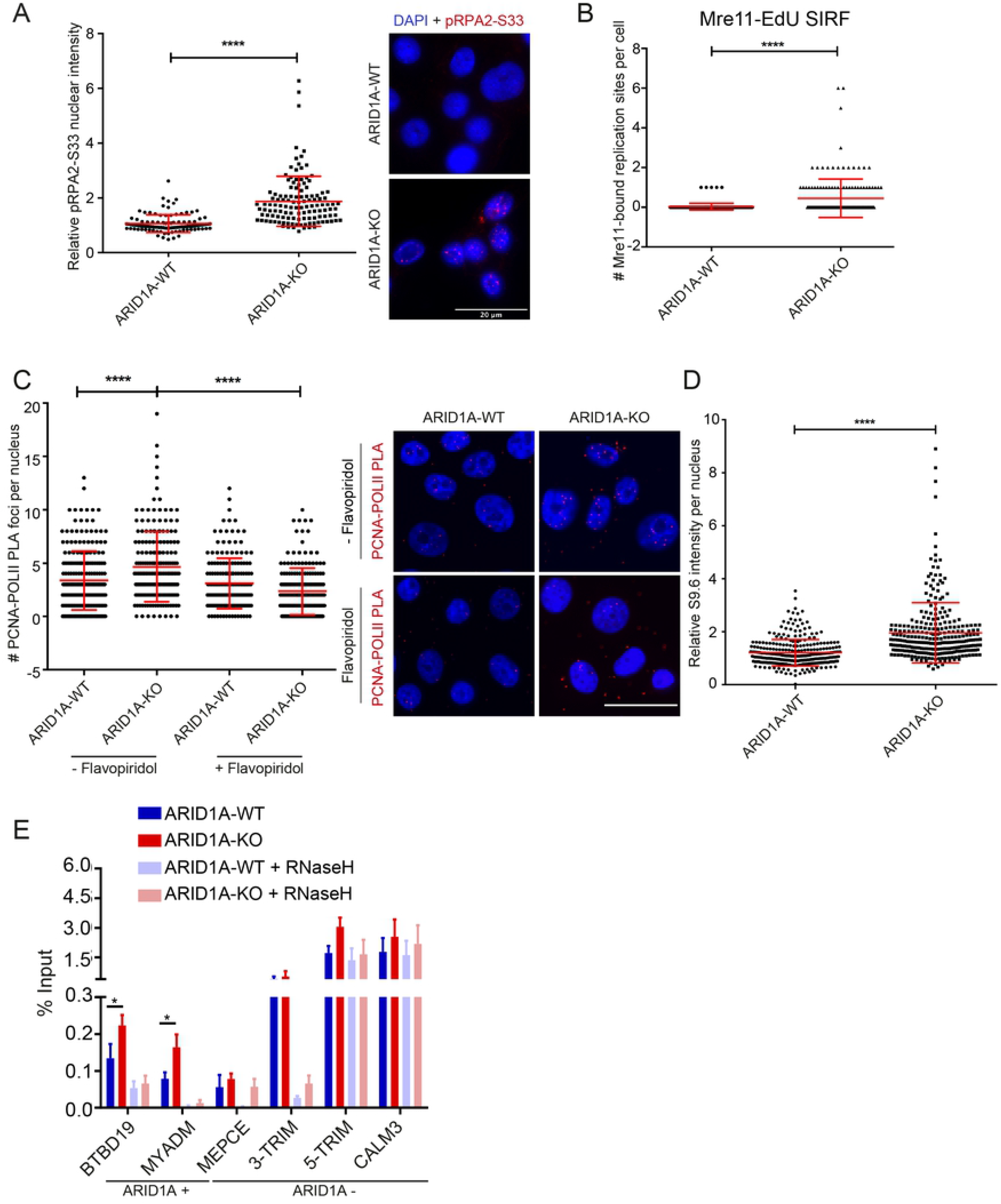
ARID1A knockout induces transcription replication conflict and R-loop accumulation. **(A)** Replication stress phenotypes in ARID1A-knockout RMG1 cells shown by relative nuclear intensity of RPA2-S33 phosphorylation in WT and ARID1A-knockout cells. Representative images, right. N=3; ****p<0.0001 by t-test; mean ± SD. **(B)** SIRF analysis of Mre11 binding to EdU-labeled nascent DNA in WT and ARID1A-KO RMG1 cells. N=3; ****p<0.0001 by t-test; mean ± SD. **(C)** Transcription-dependent regulation of transcription-replication conflicts (TRC) by ARID1A. Proximity ligation assay targeting the replisome (anti-PCNA) and RNA polymerase II (anti-RNA Pol II) is shown with quantification (left) and representative images (right). Pre-treatment with transcription inhibitor flavopiridol for 2 hours significantly reduced TRC in ARID1A-KO cells. N=3; ****p<0.0001 by ANOVA; mean ± SD. **(D)** Relative S9.6 staining intensity per nucleus in WT and ARID1A-KO RMG1 cells. N=3; ****p<0.0001 by t-test; mean ± SD. **(E)** DRIP-qPCR analysis of the indicated loci in ARID1A-WT or KO cell lines. Faded colours indicate *in vitro* RNaseH1 treatment prior to pulldown. N=3; *p<0.05 by t-test.

Given the role of the BAF complex in regulating transcriptional programs, we hypothesized that transcription-replication conflicts (TRCs) might be responsible for some of the observed replication stress. We analyzed the proximity between PCNA and RNA polymerase II as a reporter of TRCs[22,23] using a proximity ligation assay and found that ARID1A loss significantly increased such conflicts (**Figure 1C**). These signals were transcription dependent since pre-treatment with flavopiridol abolished differences between the WT and ARID1A-knockout cells (**Figure 1C**). TRCs are also often associated with increased levels of R-loop structures. To test this we used S9.6 antibody immunofluorescence which recognizes DNA:RNA hybrids and other RNA structures and found that ARID1A knockout cells had significantly higher S9.6 staining (**Figure 1D**). While the signal could be reduced in both cell lines with RNaseH or RNaseIII treatment, targeting hybrids and dsRNA respectively, increased S9.6 staining persisted in ARID1A-KO cells treated with RNaseIII supporting a true increase in R-loops (**Figure S1B**). To ensure this was not a cell-line specific effect, we stained a p53−/− derivative of HCT116 and RPE1-hTERT cells treated with siRNA targeting ARID1A and found ARID1A-depletion increased S9.6 staining (**Figure S2A** and **B**). Given concerns about the specificity of imaging with S9.6 [24], we also performed DNA:RNA immunoprecipitation (DRIP) experiments and quantitative PCR at a set of S9.6 positive loci. These included RNaseH-sensitive loci to confirm they represented DNA:RNA hybrids, and we found that the signal was increased at two of the four RNaseH sensitive sites in ARID1A-KO cells (**Figure 1E**). Importantly, when we assessed ARID1A binding at these sites in published datasets from RMG1 cells, we found that only the ARID1A+ sites BTBD19 and MYADM showed an increase in S9.6 signal when ARID1A was deleted (**Figure 1E**) [3,22,25]. Together these data show that in an RMG1 cell line model the deletion of ARID1A elicits replication stress, TRCs, and R-loop accumulation at a subset of genomic loci.

### R-loop and TRCs drive replication stress and DNA damage in ARID1A knockouts

Ectopic formation of R-loops has been linked to DNA damage, transcription-replication conflicts, replication stress and the activation of the Fanconi Anemia pathway [6,26]. To assess R-loop stabilization as a potential mechanism driving replication stress we undertook a set of suppression experiments. First, we confirmed that ectopic expression of RNaseH1-GFP strongly reduced the S9.6 staining in the ARID1A knockout cell line (**Figure 2A**). Consistent with a mechanism of R-loop induced breaks arising from replication stress, the increase in phosphorylation of RPA2-ser33 in ARID1A−/− cells was also suppressed by overexpression of RNaseH1, confirming R-loops as a driver of replication stress in these cells (**Figure 2B)**. Repair of R-loop induced replisome collisions has been reported to require members of the Fanconi Anemia pathway, such as FANCD2 [26,27]. Consistent with this model we saw significantly more FANCD2 foci by immunofluorescence in ARID1A knockout cells, and these differences were abolished when cells were ectopically expressing RNaseH1 to remove R-loops (**Figure 2C**). To confirm that these R-loops ultimately led to DNA damage we performed a neutral comet assay for DNA breaks and observed that excess damage in ARID1A-deficient cells was also suppressed by ectopic expression of RNaseH1 (**Figure 2D**). Taken together these results indicate that ectopic formation or stabilization of R-loops in cells lacking ARID1A is a driver of replication stress, activation of the Fanconi Anemia pathway and DNA damage.

**Figure 2.**
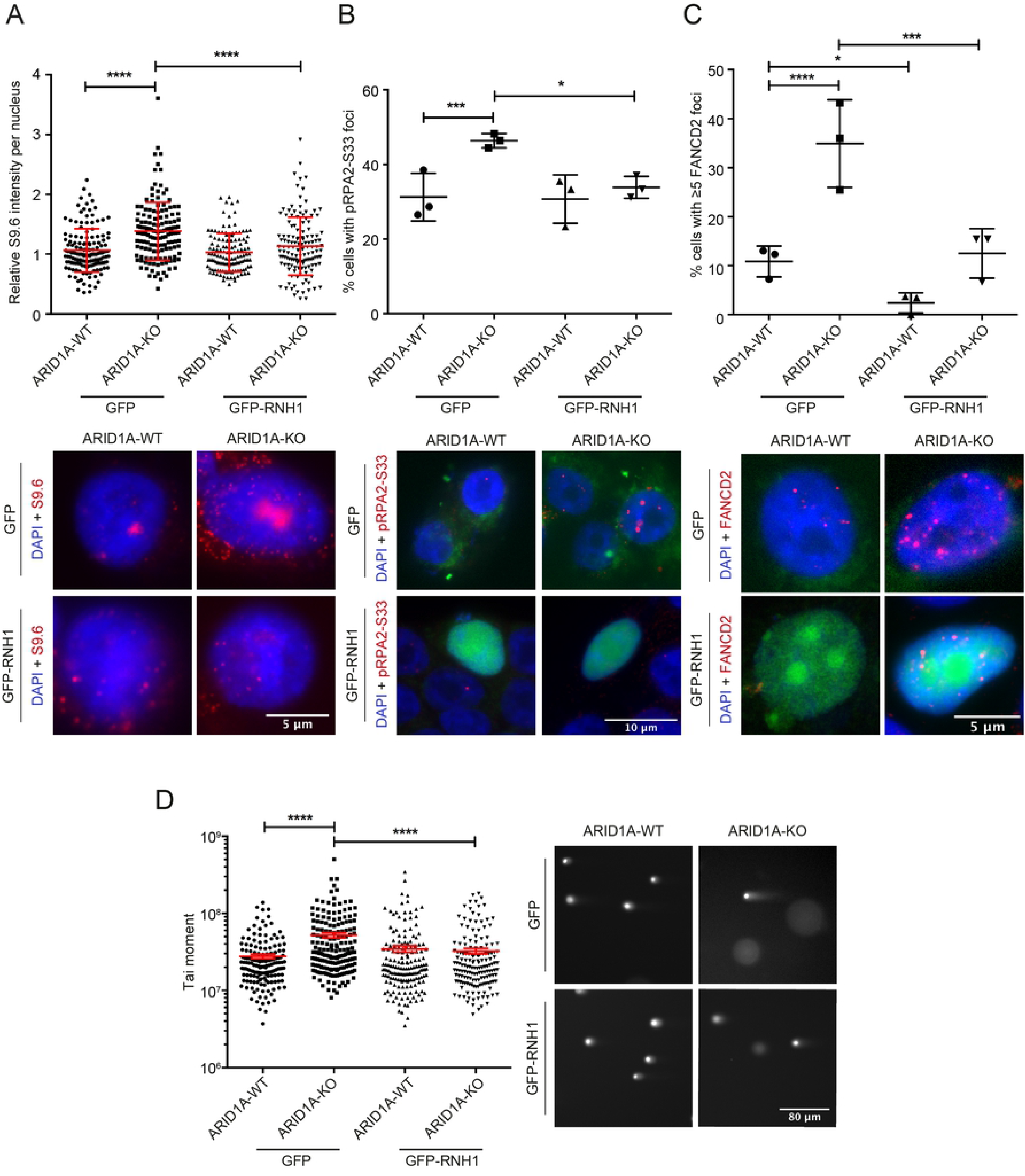
RNaseH-dependent R-loop accumulation and replication stress phenotypes in ARID1A-KO. **(A-C)** Quantification (top) and representative images (bottom) of S9.6 staining intensity per nucleus (A, left), RPA2-S33P foci (B, middle) or FANCD2 foci (C, right) in WT and ARID1A-KO RMG1 cells. Cells were transfected with either a control vector (GFP) or one expressing GFP-RNaseH1 (GFP-RNH1). For A, N=4; ****p<0.0001 by ANOVA; mean ± SD. For B, N=3; *p<0.05, ***p<0.0005 by Fisher’s exact test; mean ± SD. For C, N=3; *p<0.05, ***p<0.0005, ****p<0.0001 by Fisher’s exact test; mean ± SD. **(D)** Neutral comet assay in WT and ARID1A-KO RMG1 cells with a control GFP vector or expressing GFP-RNH1. N=3; ****p<0.0001 by ANOVA; mean ± SEM.

### ARID1a deficiency alters replication and transcription dynamics

Both decreased and increased replication fork speed have been associated with replication stress and the DNA damage response [28]. To monitor potential replication defects in our system we used DNA combing of CldU and IdU labelled fibers in ARID1A-WT or ARID1A-KO derivatives. First, we measured the ratio of CldU to IdU labelled fiber tracks and found that ARID1A-KO cells have a greater degree of asymmetry between the two labeling periods, consistent with stalled or collapsed DNA replication forks (**Figure 3A**). Surprisingly, and in spite of the presumptive increase in fork stalling, when we measured total fiber length we found that on average replication forks travel faster in ARID1A-knockouts compared to WT controls (**Figure 3B**). Since the BAF complex opens chromatin to promote transcription, one possible mechanism by which replication forks may speed up is due to reduced transcription. Indeed bulk quantification of global nascent transcription by pulse 5-ethynyl uridine (EU) incorporation showed lower overall transcription activity in ARID1A knockouts (**Figure 3C**). Consistent with this, ChIP-qPCR of RNA polymerase II also showed lower occupancy at several transcribed genes (**Supplemental Figure S3**). In addition, we found that inhibiting transcription elongation with flavopiridol, or transcription initiation with the TFIIH inhibitor triptolide also increased replication fork speed significantly in ARID1A-WT cells, but not in ARID1A-KO cells (**Figure 3D**). Global effects on transcription may explain the observed replication fork speed increases (see *Perspective* section). However, we also tested the direct association of the BAF complex with nascent DNA using SIRF for the catalytic subunit of BAF, Brg1. In this experiment, we saw a robust association in ARID1A-WT cells that was significantly reduced in ARID1A knockouts (**Figure 3E**). Therefore the BAF complex may have a previously observed but under-appreciated role in DNA replication [29]. At this point it is not clear how broad transcriptional and chromatin state changes integrate with the DNA repair, and possibly replication, functions of the BAF complex to suppress replication stress.

**Figure 3.**
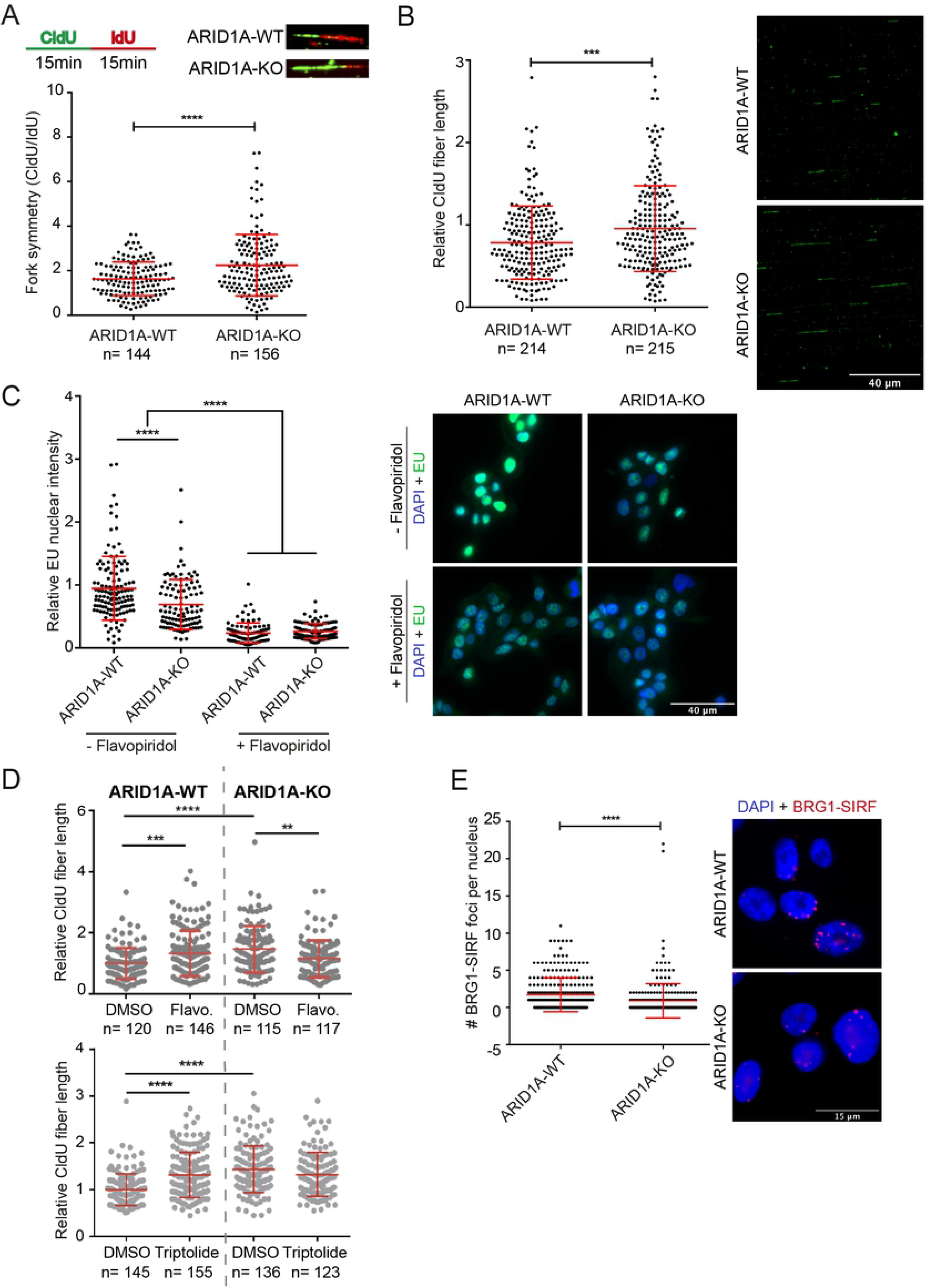
Replication and transcription dysregulation in ARID1A-KO cells. **(A)** Quantification and representative image of DNA fiber combing measurements for fork DNA replication fork symmetry in WT and ARID1A-KO RMG1 cells. Replication forks were labelled with CldU (green) for 15 minutes, followed by a 15 minute pulse of IdU (red). Replication fork symmetry was measured by calculating the length ratio of CldU/IdU tracks. N = 2; ****p<0.001 by unpaired t-test; mean ± SD. **(B)** Quantification (left) and representative images (right) of DNA fiber combing tracks used to measure DNA replication fork speed in WT and ARID1A-KO RMG1 cells. N = 3; ****p<0.0001 by unpaired T-test; mean ± SD. **(C)** Bulk quantification (left) and representative image (right) of global nascent transcription by pulse 5-ethyl uridine (EU) incorporation with or without flavopiridol treatment in WT and ARID1A-KO RMG1 cells. N = 3; ****p<0.0001 by ANOVA; mean ± SD. **(D)** Monitoring effects of transcription inhibition on replication fork speed in RMG1 ARID1A-WT and ARID1-KO cells. CldU labelling was done as in B in cells pre-treated with Flavopiridol (Flavo., Top) or Triptolide (Bottom). **p<0.01, ***p<0.001, ****p<0.0001 by unpaired t-test. For A, B and D, scored fibers numbers are shown under the axis. **(E)** Quantification (left) and representative image (right) of BRG1-SIRF foci counts in WT and ARID1A-KO RMG1 cells. N = 3; ****p<0.0001 by unpaired t-test; mean ± SD.

### Loss of ARID1A mislocalizes Topoisomerase IIα from R-loop sites

In concert with mechanisms of TRC induction involving altered transcription or replication dynamics, we also wondered if changes in the function of BAF physical interaction partners could be responsible for some of the observed phenotypes. BAF interacts with several genome stability maintenance proteins including Topoisomerase IIα (TOP2A), which has previously been implicated in regulating topological stress at TRCs [2,11]. First, we confirmed that global TOP2A inhibition with etoposide dramatically increased DNA:RNA hybrid staining with the S9.6 antibody and RPA2-ser33 phosphorylation in both ARID1A WT and knockout cell lines (**Figure 4A** and **4B**). In ARID1A knockout cells only specific genomic loci lose TOP2A localization and activity [11]. Therefore, we wanted to assess whether the association of TOP2A with R-loop prone loci was affected. First using proximity ligation assay with antibodies against TOP2A and S9.6 we saw a relative reduction in PLA foci in the ARID1A knockout cells (**Figure 4C**). Thus, despite having more R-loops overall, these data indicate that the recruitment of TOP2A to R-loop sites may be impaired in ARID1A knockouts. We considered that ARID1A loss might prevent TOP2A association with chromatin in general. However, fractionation analysis showed that TOP2A remained strongly chromatin associated in ARID1A-KO cells (**Figure S4A**). We also wondered if loss of interaction with the BAF complex was driving the change in TOP2A-S9.6 PLA signal. Surprisingly, we found through TOP2A immunoprecipitation and western blot that TOP2A was strongly associated with Brg1 and BAF155 subunits of the BAF complex regardless of ARID1A status (**Figure S4B**). While previous studies have suggested that ARID1A is a key subunit for TOP2A-BAF interaction [11], in this cell line other protein contacts, presumably with ARID1B, are sufficient to maintain a TOP2A-BAF interaction. To look at whether TOP2A recruitment to specific loci was altered we performed TOP2A ChIP analysis and found reduced TOP2A binding at R-loop prone loci (**Figure 4D**). These loci had increased DRIP signal (**Figure 1E**) showing an inverse relationship where, upon ARID1A deletion, TOP2A recruitment decreases and S9.6 binding increases. Therefore, for some loci, our data support a model in which deletion of ARID1A reduces BAF binding, fails to recruit TOP2A, and leads to accumulation of R-loops that cause transcription-replication conflicts, replication stress and ultimately DNA damage (**Figure 4E**).

**Figure 4.**
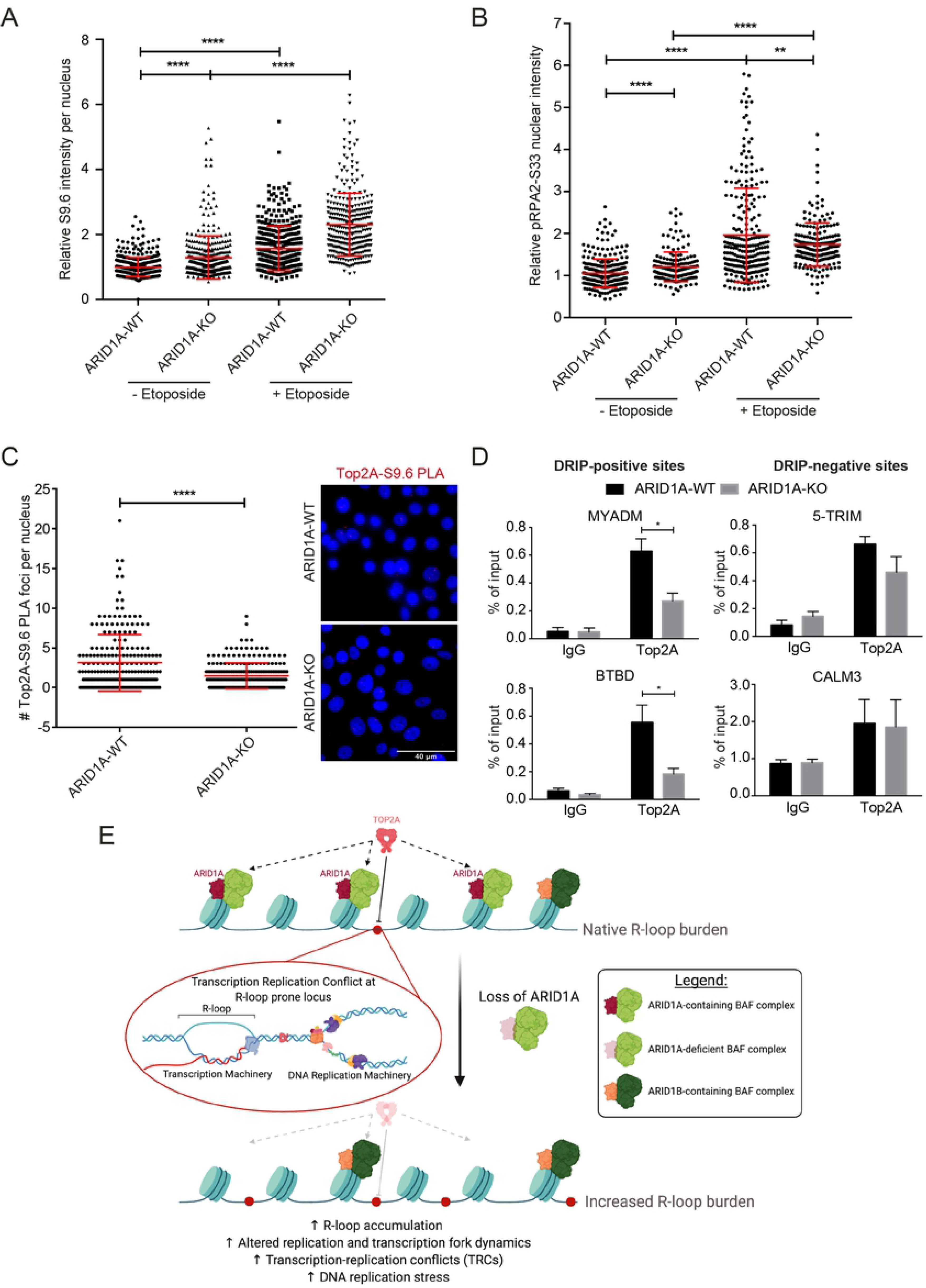
TOP2A loss from R-loop sites in ARID1A-KO cells. **(A)** Quantification of relative S9.6 nuclear intensity in WT and ARID1A-KO RMG1 cells treated with or without Etoposide (10 μM, 2 hours). Nucleolar S9.6 staining was subtracted from the total nuclear intensity. N = 3. ****p<0.0001 by ANOVA; mean ± SD. **(B)** Quantification of the relative RPA2-S33P nuclear intensity in WT and ARID1A-KO RMG1 cells treated with or without Etoposide (10 μM, 2 hours). N = 3. ****p<0.0001, **p<0.01 by ANOVA; mean ± SD. **(C)** Quantification (left) and representative images (right) of foci counts of TOP2A and S9.6 co-localization measured by proximity ligation assay. Foci represent instances when TOP2A is located in close proximity to R-loops. N= 3; ****p<0.0001 by unpaired T-test; mean ± SD. **(D)** ChIP-qPCR probing for TOP2A in WT and ARID1A-KO RMG1 cells showing loss of TOP2A binding at R-loop prone loci BTBD and MYADM, but not at RNaseH1-insensitive sites 5’-TRIM and CALM3 from Figure 1. N = 3; *p<0.05 by t-test; mean ± SEM. **(E)** Model. Under normal conditions, ARID1A-containing BAF complexes recruit TOP2A to chromatin where it can prevent the formation of TRC-inducing R-loops. Upon loss of ARID1A, TOP2A is not appropriately recruited to chromatin, resulting in increased R-loop burden, altered replication and transcription dynamics, increased incidence of TRCs and DNA replication stress. Model figure was created with BioRender.com.

### Perspective

The BAF complex has pleiotropic effects on cells due both to its transcriptional regulatory network and to an array of functions in chromatin topology, DNA repair, and replication. In cancers with disrupted BAF complexes transcriptional reprogramming likely coincides with chromatin states in the cell of origin to drive oncogenesis in specific tissues [18,30]. In addition, the BAF complex directly interacts with several candidate genome stability regulators such as ATR, BRIT1, p53, and TOP2A [11,12,31,32]. Thus, alterations in BAF can change the targeting or regulation of these proteins and potentially influence genome stability [10]. Shen et al., suggest that ARID1A recruits BAF to double strand breaks through interactions with ATR and promotes end resection [33]. The consequences of ARID1A in this setting is a homologous recombination defect and sensitivity to PARP inhibitors [33]. It is worth noting that other studies have suggested that ARID1A loss is associated with PARP inhibitor resistance [34], and therefore there remain important context-specific details to be elucidated for this approach to be successful clinically. Subsequent screening efforts identified ATR inhibition as a potentially effective synthetic lethal strategy to target ARID1A deficient tumours [19]. Our study frames these observations in a new light by directly demonstrating that ARID1A deletion increases transcription-replication conflicts and R-loop associated genome instability. Dysregulation of replication and transcriptional programs, along with altered targeting of TOP2A to R-loop prone regions underlie some of this observed stress.

Given the many roles of BAF complexes it is likely that some of the observed R-loop associated stress arises due to other mechanisms. First, BAF complexes have been implicated in DSB repair by regulating DNA end resection and Rad51 loading [13] and DNA:RNA hybrid intermediates are now known to be part of some repair reactions [35,36]. In addition, we now know that resection of stalled or reversed replication forks leads to engagement of DSB repair proteins to facilitate fork protection and restart [37,38]. Any roles for the BAF complex in this process are currently unknown. Finally, R-loops and TRCs are associated with specific chromatin states that might influence, or be influenced by, BAF complexes [39,40]. For example, recent work in yeast models has found that H3K4 methylation marks deposited by transcription are associated with reduced TRCs and genome instability by slowing incoming replication forks like ‘speed bumps’ [41]. In mammalian systems BAF is known to physically associate with H3K4me1 through its BAF45c subunit [42]. If a replication slowing mechanism proposed for this chromatin mark in yeast is conserved to human cells then our fork speed and TRC data are consistent with a chromatin-based TRC avoidance mechanism supported by ARID1A. Additional work dissecting direct and indirect roles for BAF complexes in replication dynamics will elucidate the potential coordination of chromatin states, binding partners like TOP2A, and genome stability by the BAF complex.

## ACKNOWLEDGEMENTS

We thank Robert Crouch for the RNaseH1-GFP construct, Daniel Durocher for the p53−/− RPE1-hTERT cell line, and Bert Vogelstein for the p53−/− HCT116 cell line. P.C.S. is supported by a Terry Fox Research Institute (TFRI) New Investigator award and a TFRI Program project grant. P.C.S. is a Michael Smith Foundation for Health Research Scholar and Canadian Institutes of Health Research New Investigator. L-A.F. holds a Frederick Banting and Charles Best Canada Graduate Scholarship.

## AUTHOR CONTRIBUTIONS

P.C.S. and S.T. designed the study. S.T., E.Y.C., L.F., J.P.W., S.W.M., Y.D.Z., and A.W. performed the experiments. P.C.S., S.T., L.F and E.Y.C. wrote the manuscript. Y.W. and D.G.H. provided key reagents.

## MATERIALS AND METHODS

### Cell culture and transfection

RMG1 cells were cultivated in RPMI-1640 medium (Stemcell technologies) supplemented with 10% fetal bovine serum (Life Technologies) in 5% CO_2_ at 37°C. ARID1A was knocked out in RMG1 cells using CRISPR/cas9 technology with gRNA targeting exon 2 of ARID1A gene (5’-CTTGCTGCGGTCCTGACGGAGG-3’). Targeted sequencing with Illumina MiSeq confirmed homozygous deletion (c.1615 del C) in RMG1_AC14 ARID1A knockout single clone. For RNA interference, cells were transfected with siGENOME-SMARTpool siRNAs from Dharmacon (Non-targeting siRNA Pool #1 as si-Cont and si-ARID1A). Transfections were done with Dharmafect1 transfection reagent (Dharmacon) according to manufacturer’s protocol and harvested 48 hours after the siRNA administration. For experiments with overexpression of GFP or nuclear-targeting GFP-RNaseH1 (gift from R. Crouch), transfections were performed with Lipofectamine 3000 (Invitrogen) according to manufacturer’s instructions 24 hours after the siRNA transfections.

### Immunofluorescence

For all immunofluorescence experiments, cells were grown on coverslips overnight before fixing. For experiments with GFP or GFP-RNH1 overexpression, plasmids were transfected 24hours post-seeding and were fixed 24-48 hours post transfection. For S9.6 staining, cells were fixed with ice-cold methanol for 10 minutes and permeabilized with ice-cold acetone for 1 minute. For all other stainings, cells were fixed with 4% paraformaldehyde for 10 minutes and permeabilized with 0.2% Triton X-100 for 10 minutes on ice. After permeabilization, cells were washed with PBS and blocked in 3%BSA, 0.1% Tween 20 in 4X saline sodium citrate buffer (SSC) for 1 hour at room temperature. Cells were then incubated with primary antibody for 1 hour at room temperature or overnight at 4°C. Following PBS wash, cells were then incubated with Alexa-Fluoro-488 or 568-conjugated secondary antibodies for 1 hour at room temperature, washed with PBS for 3 times, and stained with DAPI before mounting and imagining on LeicaDM18 microscope at 100X. ImageJ was used for image processing and quantification [43]. For the quantification of S9.6 intensity in RNaseIII and etoposide treated cells, cells were co-stained with anti-nucleolin antibody (abcam, ab22758, 1:1000) to mask out nucleolin-stained regions in order to quantify nuclear S9.6 signal only, outside of nucleoli, since nucleoli are hotspots of R-loop staining. For *in vitro* RNaseH or RNaseIII treatment, cells were treated with RNaseH (New England Biolabs) for 2 hours or ShortCut RNaseIII (New England Biolabs) for 20 minutes at 37°C after permeabilization before blocking.

### Proximity ligation assay and SIRF

PLA experiments were performed using the Duolink PLA kit (Millipore Sigma). For all PLA experiments, cells were grown on coverslips overnight before fixing. For S9.6 staining, cells were fixed with ice-cold methanol for 10 minutes and permeabilized with ice-cold acetone for 1 minute. For all other staining, cells were fixed with 4% paraformaldehyde for 10 minutes and permeabilized with 0.2% Triton X-100 for 10 minutes on ice. After permeabilization, cells were washed with PBS and blocked 1 hour at RT in Duolink Blocking Solution. Cells were then incubated with primary antibodies diluted in Duolink Antibody Diluent for 1 hour at room temperature or overnight at 4°C. Cells were washed twice (5 min) in Wash Buffer A, after which they were incubated 1 hour at 37°C with PLA probe mix (15ul/ cover slip, 1:4 PLA probe in PLA Antibody Diluent). Cells were washed (2x 5 min) with PLA Wash Buffer A, after which they were incubated 30 minutes at 37°C with PLA ligation mix (15ul/cover slip, 1:40 PLA ligase 40X in 1:5 ligation buffer 5X in Ultra H2O). Cells were washed (2x 2 min) with PLA Wash Buffer A and incubated 100 minutes at 37°C with PLA Amplification mix (15ul/slide, 1:80 polymerase solution in 1:5 amplification stock in ultra H2O). The cells were washed (2x 10 min) in PLA Wash Buffer B. After an additional wash (1 minute) in PLA Wash Buffer B 0.01X, the slides were mounted in Duolink Mounting Media with DAPI. Imaging was performed on a Leica Dmi8 microscope at 100X. ImageJ was used for image processing and quantification. SIRF (in situ protein interactions at nascent and stalled replication forks) was carried out exactly as described previously [20]. To inhibit transcription in control experiments, cells were pre-treated with DMSO or 0.8μM flavopiridol for 2 hours before fixation.

### Comet assay

Neutral comet assay was performed as previously described[22] using the CometAssay Reagent Kit for Single Cell Gel Electrophoresis Assay (Trevigen) in accordance with the manufacturer’s instructions.

### DNA fiber assay

DNA fiber assay was performed as previously described (22) with some modification. Cells were untreated, treated with DMSO, or treated with 1mM tripolide (Sigma) for 2 hours and then labeled with CldU only for 15mins, or were pulse labeled for 15 minutes with 30μM CldU (Sigma), washed twice with PBS, and then pulse labeled with 250μM IdU (Sigma) for 15min. Cells were then collected and scraped into ice-cold PBS and genomic DNA was extracted with CombHeliX DNA Extraction kit (Genomic Vision) in accordance with the manufacturer’s instructions. DNA fibers were stretched on vinyl silane-treated glass coverslips (CombiCoverslips) (Genomic vision) with an automated Molecular Combing System (Genomic Vision). After combing, the stretched DNA fibers were dehydrated in 37°C for 2 hours, fixed with MeOH:Acetic acid (3:1) for 10 minutes, denatured with 2.5 M HCl for 1 hour, and blocked with 5% BSA in PBST for 30 minutes. IdU and CldU were then detected with the following primary antibodies in blocking solution for 1 hour at room temperature: mouse anti-BrdU (B44) (1:25) (BD) for IdU and rat anti-BrdU [Bu1/75 (ICR1)] (1:100) (abcam) for CldU. After PBS wash, fibers were than incubated with secondary antibodies anti-Rat-Alexa488 (1:50) (Invitrogen) and anti-mouse-Alexa 568 (1:50) (Life Technologies) for 1 hour at room temperature. DNA fibers were analyzed on LeicaDM18 microscope at 100X and ImageJ was used to measure fiber length.

### DRIP and ChIP-qPCR

Cells were crosslinked in 1% formaldehyde for 10 minutes before quenching with glycine for 5 minutes at room temperature, and then lysed in ChIP lysis buffer (50mM HEPES-KOH at pH 7.5, 140 mM NaCl, 1 mM EDTA at pH 8, 1% Triton X-100, 0.1% Na-Deoxycholate, 1% SDS) and rotated for 1 hour at 4 °C. DNA were sonicated on a Q Sonica Sonicator Q700 for 8 minutes (30 sec ON, 30 sec OFF) to generate fragments of 200-500bp. For DRIP, the chromatin preps were treated with 20 mg/mL Proteinase K (Thermo Fisher Scientific) at 65°C overnight and total DNA was purified by phenol/chloroform purification method. Protein A magnetic beads (Bio-rad) were first pre-blocked with PBS/EDTA containing 0.5% BSA and then incubated with S9.6 antibody (1:200, clone S9.6, MABE1095, Millipore) in IP buffer (50 mM Hepes/KOH at pH 7.5; 0.14 M NaCl; 5 mM EDTA; 1% Triton X-100; 0.1% Na-Deoxycholate, ddH2O) at 4°C for 4 hour with rotation. DNA was then added to the mixture and gently rotated at 4°C overnight. Beads were recovered and washed successively with low salt buffer (50mMHepes/KOH pH 7.5, 0.14 M NaCl, 5 mM EDTA pH 8, 1% Triton X-100, 0.1% Na-Deoxycholate), high salt buffer (50 mM Hepes/KOH pH 7.5, 0.5 M NaCl, 5 mM EDTA pH 8, 1% Triton X-100, 0.1% Na-Deoxycholate), wash buffer (10 mM Tris-HCl pH 8, 0.25 M LiCl, 0.5% NP-40, 0.5% Na-Deoxycholate, 1 mM EDTA pH 8), and TE buffer (100 mM Tris-HCl pH 8, 10 mM EDTA pH 8) at 4°C, two times. Elution was performed with elution buffer (50mMTris-HCl pH 8, 10mMEDTA, 1% SDS) for 15 minutes at 65°C. After purification with PCR Cleanup kit (Sigma-Aldrich), nucleic acids were eluted in 100 μL of elution buffer (5 mM Tris-HCl pH 8.5) and analyzed by quantitative real-time PCR (qPCR). For ChIP, DNA and antibody were incubated in IP buffer overnight with rolling at 4°C. Antibodies used were RNA polymerase II (abcam), BRG1 (Santa Cruz), TOP2A (Santa Cruz). Antibody-bound DNA was recovered using the Protein A or Protein G magnetic beads (Bio-Rad), washed similarly as DRIP samples and treated with Proteinase K and RNAse after elution. Then, antibody-bound DNA was purified with PCR Cleanup kit and analyzed by qPCR. qPCR was performed with Fast SYBR Green Master (ABI) on AB Step One Plus real-time PCR machine (Applied Biosystem). Primer sequences are listed in table 1. qPCR results were analyzed using the comparative CT method. The RNA-DNA hybrid and ChIP DNA enrichments were calculated based on the IP/Input ratio. ARID1A positive sites were determined from GEO dataset GSM3392689 [44].

### Nascent RNA labeling

Cells were seeded on coverslips and grown overnight. The next day cells were incubated with 0.5 mM EU for 1 hour. After EU labeling cells were washed with PBS twice and then fixed with 4%PFA and permeabilized with 0.25% TritonX-100 in PBS. EU incorporation was measured with Click-iT^®^ RNA Imaging Kits (Invitrogen) using Alexa Fluor 488 dye according to manufacturer’s instruction. Slides were stained with DAPI before mounting and imaging on a LeicaDM18 microscope at 100X. ImageJ was used for image processing and quantification. For negative control, cells were first treated with 0.8μM flavopiridol for 2 hours before EU labeling.

### Immunoprecipitation (IP) and Western blotting

Whole cell lysates were prepared with RIPA buffer containing protease inhibitor (Sigma) and phosphatase inhibitor (Roche Applied Science) cocktail tablets. For nuclear plasma and chromatin fraction proteins extraction, cells were lysed by 10 mM HEPES pH 7.4, 10 mM KCl, 0.05% NP-40 and incubated 20 min on ice then centrifuged and removed supernatant to keep nuclei pellet. The nuclei pellet was lysed with 10 mM Tris-HCl pH 7.4, 0.2 mM MgCl2, 1% triton X-100 and incubated 15 min on ice then centrifuged at 14,000rpm at 4°C. The supernatant was nuclear plasma proteins and the pellet was lysed with HCl 0.2N and incubate 20 min on ice then centrifuged at 14,000 rpm at 4°C, 10 min. Then added the same volume of 1M Tris-HCl pH 8 to neutralize supernatant chromatin proteins. For IP, cells were lysed by 10 mM HEPES pH 7.4, 10 mM KCl, 0.05% NP-40 and incubated 20 min on ice then centrifuged and removed supernatant to keep nuclei pellet. The total nuclear protein was lysed with CHAPS buffer containing protease inhibitor (Sigma) and phosphatase inhibitor (Roche Applied Science) cocktail tablets. The protein concentration was determined by Bio-Rad Protein assay (Bio-Rad). Protein lysates were pre-clear in Protein G magnetic beads (Bio-rad) at 4°C for 2 hours with rotation. Protein G magnetic beads were incubated with TOP2A antibody (Santa Cruz) in CHAPS buffer at 4°C for 2 hours with rotation. Protein extract was then added to the mixture and gently rotated at 4°C overnight then beads were washed 4 times with CHAPS buffer and then boiled in Laemmli Sample Buffer (Bio-Rad) for 10 min. Equivalent amounts of protein or IP pulldown samples were resolved by SDS-PAGE and transferred to polyvinylidene fluoride (PVDF) microporous membrane (Millipore), blocked with 5% skim milk in TBS containing 0.1% Tween 20 (TBST), and membranes were probed with the following antibodies: ARID1A, Top2a, Lamin B1, BRG1, (1:250) (Santa Cruz), BAF155 (1:1000) (cell signaling) and GAPDH (1:5000) (ThermoFisher Scientific). Secondary antibodies were conjugated to Horseradish Peroxidase and peroxidase activity was visualized using Chemiluminescent HRP substrate (Thermo Scientific).

